# Mitochondrial ATP-dependent K^+^ channels downregulate ionotropic glutamate receptors during hypoxia in retinal horizontal cells of goldfish

**DOI:** 10.64898/2026.09.28.754954

**Authors:** Mohamed Ramadan, Christian A. Rosales, Jeffrey C. Smith, Michael G. Jonz

**Author notes:** Author for correspondence: Michael G. Jonz, Ph.D., Department of Biology, University of Ottawa, 30 Marie Curie Pvt., Ottawa, ON K1N 6N5, Canada.

## Abstract

Retinal neurons rely on continuous oxygen supply to maintain function, yet some species, like goldfish, withstand prolonged hypoxia due to adaptations that remain unclear. Horizontal cells (HCs) are inhibitory interneurons of the retina that express ionotropic glutamate receptors (iGluRs) and secrete γ-aminobutyric acid (GABA), allowing them to shape visual signalling by feeding back to photoreceptors. We investigated whether iGluRs in dissociated goldfish HCs are suppressed during hypoxia, and whether this process is mediated by oxygen-sensitive activation of mitochondrial ATP-sensitive K^+^ (mK_ATP_) channels. Through perforated patch-clamp electrophysiology, we determined that 20 min of hypoxia reduced peak inward current density associated with glutamate application by ∼25% (P = 0.0059). When mK_ATP_ channels were selectively blocked with 100 µM glibenclamide, hypoxic suppression of iGluRs was abolished. Inhibiting mitochondrial Ca^2+^ uptake via the mitochondrial Ca^2+^ uniporter with ruthenium red, or blocking Ca^2+^ release through ryanodine receptors in the endoplasmic reticulum, also abolished the hypoxia-induced suppression. Similarly, inhibition of calmodulin with calmidazolium, or of protein phosphatase 1 and 2A (PP1/PP2A) activity with calyculin A, prevented the hypoxia-induced suppression of iGluRs. Taken together, these results indicate that mK_ATP_ activation is required to mediate an intracellular Ca^2+^ signal that downregulates iGluR activity during hypoxia. Through the reduction of iGluR activity, HCs may lower their metabolic demand, allowing them to maintain cell function during hypoxia. Additionally, GABA release from HCs, as measured by mass spectrometry, was not affected by hypoxia, and we propose that downregulation of iGluR may account for hypoxic suppression of the retina in hypoxia-tolerant organisms.

**SUMMARY STATEMENT:** Mitochondrial ATP-dependent K^+^ channels contribute to hypoxia-induced suppression of ionotropic glutamate receptor activity in goldfish retinal horizontal cells.

## INTRODUCTION

The vertebrate retina has a high and continuous demand for energy to maintain ion gradients and synaptic transmission. Horizontal cells (HCs) are inhibitory interneurons of the outer retina that receive glutamatergic input from photoreceptors and provide feedback that contributes to visual contrast and colour opponency (Thoreson & Mangel, 2012; Dowling, 1987). In darkness, photoreceptors are depolarized and tonically release glutamate onto HCs. This glutamatergic signal provides the primary excitatory signal from photoreceptors to downstream neurons in the retina (Massey, 1990; Cervetto & Piccolino, 1974; Thoreson & Witkovsky, 1999). Activation of ionotropic glutamate receptors (iGluRs) depolarizes the HC membrane, opening voltage-gated Ca^2+^ channels and increasing intracellular Ca^2+^ concentration ([Ca^2+^]_i_) (Linn and Christensen, 1992; Tachibana, 1983). Maintaining this activity requires ATP-dependent pumps to restore ionic gradients and keep [Ca^2+^]_i_ within a physiological range (Country, 2017; Erecińska & Silver, 1989; Garret & Ross 1985). When oxygen availability becomes insufficient to meet cellular demand (i.e. hypoxia), ATP production declines and these homeostatic processes are disrupted. Uncontrollable membrane depolarization, reduced glutamate uptake and excessive glutamate release can then prolong iGluR activation and increase [Ca^2+^]_i_ to toxic levels, resulting in excitotoxic cell death (Szydlowska & Tymianski, 2010; Orrenius & Nicotera, 1994; Orrenius et al., 2003; Choi, 1992; Lucas & Newhouse, 1957; Verkhratsky, 2007).

Some vertebrates have evolved mechanisms that prevent this response and permit neuronal survival in the central nervous system, including the retina, during prolonged hypoxia or complete anoxia (Walker & Johansen, 1977). Painted turtles (*Chrysemys picta*), crucian carp (*Carassius carassius*) and common goldfish (*Carassius auratus*) can reversibly reduce neural activity and ATP consumption when oxygen is limited (Hylland et al., 1997; Johansson et al., 1997,1995; Nilsson & Lutz, 2004; Bickler & Buck, 2007; Wilkie et al., 2008; Pamenter et al., 2011; Country & Jonz, 2021, 2022). This suppression of neural activity during oxygen deprivation is consistent with the channel arrest hypothesis proposed by Hochachka (1986), which suggests that cells reduce membrane ion permeability and ATP-dependent ion pumping to preserve electrochemical gradients while lowering energetic demand during metabolic stress.

Mitochondrial ATP-sensitive K^+^ (mK_ATP_) channels may provide a cellular mechanism for initiating channel arrest during oxygen deprivation. As ATP levels decline, activation of mK_ATP_ channels produces a mild mitochondrial depolarization and initiate protective intracellular signalling (Inoue et al., 1991; Country & Jonz, 2021; Buck & Pamenter, 2018). In turtle cortical neurons mK_ATP_ activation limits excitatory glutamate receptor activity during anoxia (Pamenter et al., 2008). In HCs isolated from hypoxia-intolerant rainbow trout, pharmacological activation of mK_ATP_ prevented a harmful rise in [Ca^2+^]_i_ (Country & Jonz, 2021). mK_ATP_ channels also contribute to ischemic preconditioning in the mammalian retina, in which a brief nonlethal ischemic exposure protects retinal tissue against a later, more severe insult (Roth et al., 2006).

Recent studies suggest that goldfish HCs may engage a similar channel arrest pathway to limit membrane ion flux and reduce energetic demand during hypoxia, although how this response relates to iGluR activity has not been established. In the absence of glutamate, goldfish HCs maintain stable [Ca^2+^]_i_ during prolonged hypoxia through an mK_ATP_-dependent mechanism (Country & Jonz, 2021). Paradoxically, when HCs are activated by glutamate during hypoxia, a modest increase in [Ca^2+^]_i_ is observed. This response is abolished by the inhibition of the mitochondrial Ca^2+^ uniporter or ryanodine receptors, indicating that mitochondrial Ca^2+^ handling and release from the endoplasmic reticulum contribute to the signal (Nagy-Watson & Jonz, 2025). How this glutamate-dependent Ca^2+^ signal in inhibitory HCs may lead to retinal suppression during oxygen deprivation (Johansson et al., 1997) remains unclear. Establishing this relationship is important for defining the cellular mechanisms that underlie hypoxia tolerance in goldfish retina.

One possibility is that the increase in [Ca^2+^]_i_ recruits intracellular signalling mechanisms that supress iGluR activity. In turtle cortical neurons, inhibition of calmodulin or protein phosphatases 1 and 2A (PP1/PP2A) prevents the anoxia induced reduction in NMDA receptor activity (Shin et al., 2005; Bickler et al., 2000). These findings identify calmodulin and PP1/PP2A as potential components linking Ca^2+^ signalling to glutamate receptor suppression. A similar mechanism in goldfish HCs could reduce ion flux and the energetic cost of maintaining ionic gradients. Whether these signalling components contribute to hypoxia-induced suppression of iGluRs in HCs remains unknown.

A second possibility that may explain suppression of retinal activity is that the Ca^2+^ signal promotes release of γ-aminobutyric acid (GABA) from HCs and thereby supresses retinal activity. The reversible suppression of visual responses in crucian carp during anoxia suggest a potential role for HCs in retinal suppression. Goldfish HCs store and release GABA, a process that is partly Ca^2+^-dependent (Ayoub and Lam, 1984,1985; Barnes et al., 2020; Marc et al., 1978). The rise in [Ca^2+^]_i_ observed during hypoxia in glutamate-stimulated HCs therefore raises the possibility of increased GABA release during oxygen deprivation. GABAergic signaling has been shown to contribute to neural suppression in the anoxic turtle and crucian carp brain (Nilsson and Lutz, 1991; Nilsson, 1992; Pamenter et al., 2011), supporting the possibility of a similar inhibitory response in the retina. However, whether hypoxia increases GABA release from goldfish HCs remains unclear.

The present study examined whether the hypoxia-induced increase in [Ca^2+^]_i_ previously observed in goldfish HCs contributes to inhibitory signalling within the retina. Specifically, we tested whether this Ca^2+^ response suppresses iGluR activity and/or promotes GABA release during hypoxia. These experiments were designed to clarify how Ca^2+^ signalling contributes to the cellular mechanisms that support retinal function during oxygen deprivation.

## MATERIALS AND METHODS

### Animals

Animal care procedures were approved by the University of Ottawa Animal Care and Veterinary Services (protocol BL-3677) and followed the guidelines set by the Canadian Council on Animal Care. Adult common goldfish (*Carassius auratus*, Linnaeus 1758) weighing 7–84 g were obtained from Mirdo Importations Canada (Montreal, QC, Canada) and housed in the University of Ottawa Laboratory for the Physiology and Genetics of Aquatic Organisms. Fish were kept in 170 L tanks with a continuous flow of fresh, aerated and dechloraminated water and held at 18°C. A 12h light:12h dark photoperiod was maintained. Prior to experimentation, goldfish were dark adapted for at least 1 h before being euthanized. All datasets presented in this study were obtained from recordings of cells collected from both male and female individuals of different ages.

### Isolated cell preparation

Horizontal cells were isolated using a protocol modified from Jonz and Barnes (2007). Goldfish were euthanized by concussion, rapidly decapitated and pithed. The eyes were enucleated and placed in ice-cold, Ca²⁺-free Ringer’s solution containing the following (in mM): 120 NaCl, 2.6 KCl, 1 NaHCO₃, 0.5 NaH₂PO₄, 1 sodium pyruvate, 4 HEPES and 16 glucose, adjusted to pH 7.8 with NaOH. The optic nerve was cut, and the sclera and lens were removed to provide access to the retina. Retinas were extracted from the eyecups and transferred to a solution consisting of 70% Leibovitz’s L-15 medium and 30% Ca²⁺-free Ringer’s solution, hereafter referred to as L-15. Retinas were incubated in L-15 containing 100 U mL⁻¹ hyaluronidase (Cat. No. H3506, Sigma, Mississauga, ON, Canada) for 20 min at room temperature to remove excess vitreous material.

### Electrophysiology

Patch-clamp recordings were performed using the perforated-patch configuration. Electrodes were pulled on the day of recording from borosilicate capillary glass (B150-86-10, Sutter Instruments, Novato, CA, USA) using a P-1000 horizontal micropipette puller (Sutter Instruments). Electrodes had resistances of 6–10 MΩ. Electrode tips were filled with intracellular solution containing (in mM): 10 NaCl, 120 KCl, 2 MgCl_2_, and 10 HEPES with pH adjusted to 7.4 with KOH. Electrodes were then backfilled with a solution of amphotericin B at a working concentration of 62.5 µg/mL. Only cells with a stable membrane seal of >1 GΩ were used. Isolated cells were plated into a 35 mm Petri dish fitted with a perfusion chamber. Throughout recordings, cells were continuously superfused at a rate of at ∼1 mL min^−1^ using a gravity-driven perfusion system. A peristaltic pump simultaneously removed solution at the same flow rate to prevent fluctuation of fluid level in the chamber.

All protocols were preformed with pClamp 7 software, an Axopatch 1D amplifier, and a Digidata 1440A (Axon Instruments, Sunnyvale, CA, USA). All cells were clamped at a holding potential of –60 mV. Macroscopic membrane currents were recorded using voltage-ramp protocols in which the membrane potential (*V_m_*) was gradually depolarized from –120 mV to +100 mV over 1 s. Additionally, gap-free recordings were also conducted to monitor spontaneous current activity over extended periods of time (25–45 min). Throughout these recordings, various drugs were applied through perfusion and their effects on membrane current were recorded. Membrane capacitance (*C_m_*) was measured with pClamp 7.

### Experimental procedures and solutions

After plating, cells were given 10-15 min to contact the bottom of the dish before any patch-clamp experiments began. Cells were washed with extracellular solution (ECS) in mM: 120 NaCl, 5 KCl, 2.5 CaCl_2_, 2 MgCl_2_, 10 HEPES, and 10 glucose with pH adjusted to 7.8 with NaOH, for a few minutes before recording to remove any debris. In all experiments HCs were treated with normoxic ECS for a few minutes to establish a stable baseline recording. Hypoxic exposure lasted no longer than 2 min during voltage-ramp protocols, and 20 min or longer during gap-free recordings.

Hypoxic solutions were prepared by bubbling solution reservoirs with 100% N_2_ for a minimum of 20 min prior to experiments, resulting in a recording solution with a PO_2_ of approximately 25 mmHg (Nagy-Watson & Jonz, 2025). Control solutions were bubbled with air for the same duration. For voltage-ramp recordings, HCs were treated with glutamate for 2 min at three time points: (1) in normoxic conditions as a control, (2) during hypoxic exposure, and (3) following a washout period. Similar experiments were conducted with normoxic ECS to serve as a sham control. For gap-free recordings, HCs received sequential 2-min glutamate applications separated by a washout period in ECS under continuous normoxia or hypoxia. The total time of these experiments was 25 min. Glutamate (100 μM; Cat. No. 49621, Sigma), and ruthenium red (400 nM; Cat. No. 1439, Tocris) were dissolved in ECS. Ryanodine (20 μM; Cat. No. 1329, Tocris), glibenclamide (100 μM; Cat. No. G0639, Sigma), calmidazolium chloride (15 μM; Cat. No. 14442, Cayman Chemical Company) and calyculin A (10 nM; Cat. No. 208851, Sigma) were first dissolved in dimethyl sulfoxide (DMSO, Cat. No. D8418), which never surpassed a final concentration of 0.25% v/v, before dilution in ECS. At this concentration, DMSO had no effect on membrane currents. The concentrations of all solutions used are summarized in Table 1.

**Table 1.** Composition of extracellular solutions (ECS) used in the present study. pH was adjusted to 7.8 (± 0.01) with NaOH for all solutions. Column labels across the top show that the first column is control ECS, whereas the remaining columns are ECS with the added drugs indicated. Chemical composition of each solution is indicated in the leftmost column.

| **Solutions and chemical concentrations (in mM)** |  |  |  |  |  |  |  |  |
| --- | --- | --- | --- | --- | --- | --- | --- | --- |
|  | ECS | Calyculin A | Calmidazolium | Glutamate | Glibenclamide | Glibenclamide + Glutamate | Ruthenium Red + Glutamate | Ryanodine + Glutamate |
| **Chemical** |  |  |  |  |  |  |  |  |
| NaCl | 120 | 120 | 120 | 120 | 120 | 120 | 120 | 120 |
| KCl | 5 | 5 | 5 | 5 | 5 | 5 | 5 | 5 |
| CaCl<sub>2</sub> | 2.5 | 2.5 | 2.5 | 2.5 | 2.5 | 2.5 | 2.5 | 2.5 |
| MgCl<sub>2</sub> | 2 | 2 | 2 | 2 | 2 | 2 | 2 | 2 |
| HEPES | 10 | 10 | 10 | 10 | 10 | 10 | 10 | 10 |
| Glucose | 10 | 10 | 10 | 10 | 10 | 10 | 10 | 10 |
| Glutamate | - | - | - | 0.1 | - | 0.1 | 0.1 | 0.1 |
| Glibenclamide<sup>1</sup> | - | - | - | - | 0.1 | 0.1 | - | - |
| Ruthenium Red | - | - | - | - | - | - | $4 \times 10^{-4}$ | - |
| Ryanodine<sup>1</sup> | - | - | - | - | - | - | - | 0.2 |
| Calmidazolium<sup>1</sup> | - | - | 0.015 | - | - | - | - | - |
| Calyculin A<sup>1</sup> | - | $1 \times 10^{-5}$ | - | - | - | - | - | - |

**1.** Dissolved in DMSO to a final concentration of 0.25% v/v.

### GABA release experimental procedures and solutions

Retinas were removed and treated with enzymes for dissociation (as described above), then rinsed an additional three times in ECS. The experimental design ensured that each goldfish contributed to three equal samples across treatment groups to minimize variability. These samples were then assigned to one or more of the following four treatment conditions: 1) control (ECS), 2) glutamate, 3) hypoxia, or 4) hypoxia + glutamate. The specific combination of treatments varied depending on the experimental setup. In some cases, samples received different treatments. In other cases, one or more samples received the same treatment. For example, one fish might have two samples treated with hypoxia and glutamate and one with glutamate alone. Cells were exogenously treated with glutamate, which was diluted in ECS to a final concentration of 100 μM (Nagy-Watson & Jonz, 2025).

The data presented from GABA analysis are derived from 22 dishes of HCs, with each dish representing an individual sample. All 22 samples were collected from 8 goldfish, with each goldfish (2 retinas) yielding 3 dishes. The cell treatment suspensions were transferred to 35 mm culture dishes and placed in an atmosphere-controlled incubator (Forma 3130, ThermoFisher Scientific, Ottawa, ON, Canada) at 22-24°C. Experimental samples were subjected to hypoxic conditions (P_O2_ = 15 mmHg) for 1 h, while control samples remained in normoxia for the same duration. After incubation, cell suspensions from each dish were moved into 15 mL Falcon tubes and centrifuged at 1100 RPM for 5 min, or until a pellet formed. The supernatant was collected, passed through a 0.2 μm syringe filter, and transferred into a microcentrifuge tube. Finally, the samples were stored at −20°C and sent to the Carleton Mass Spectrometry Centre at Carleton University to test for the presence of GABA using liquid chromatography-mass spectrometry (LC-MS) analysis. Samples were dried to completeness using a stream of N_2_ and reconstituted in an aqueous caffeine diluent internal standard (IS) mixture (0.5 µM) prior to LC-MS analysis. Standard samples were reconstituted in the same volume of IS diluent as the original sample aliquot amount, whereas saline culture media and goldfish retina supernatant samples were diluted 5-fold to mitigate matrix effects.

### Materials for chemical analysis, reversed-phase liquid chromatography, and ESI-multiple reaction monitoring (MRM) analysis

We developed a procedure for the detection of GABA in the supernatant samples using LC-MS (Rosales et al., 2025). GABA and caffeine were purchased from Sigma-Aldrich (St. Louis, MO, USA). LC-MS grade water with 0.1% formic acid and acetonitrile with 0.1% formic acid were purchased from Fisher Scientific (Hampton, NH, USA). A sample volume of 20 µl was injected into a Dionex Ultimate 3000 HPLC (Thermo Fisher Scientific, Waltham, MA, USA) and separated on an Agilent Eclipse plus C18 3.5 µm column (4.6 × 100 mm) at room temperature.

At a constant flow rate of 0.6 mL/min, analytes were eluted using eluent A (water + 0.1% formic acid) and eluent B (acetonitrile + 0.1% formic acid) over a 10.5 min gradient [0.0-1.0 min, 0.5% (B); 5.0 min, 50% (B); 5.1-8.0 min, 100% (B); 8.1-10.5 min, 0.5% (B)]. The HPLC eluent was coupled to a 4000 QTrap hybrid triple quadrupole linear ion trap mass spectrometer (Sciex, Framingham, MA, USA), analyzing all samples in positive ion mode. The Turbo Spray ion source parameters were as follows: curtain gas of 20, high collision gas, 5000 V ionization voltage, temperature of 375°C, ion source gas 1 of 30, and ion source gas 2 of 10. All samples were analyzed using optimized MRM transitions at 20 Hz (50 ms), with each analyte having a transition pair for semi-quantitation and validatory identification (Table 2).

**Table 2.** Optimized MRM transition pairs for GABA and the internal standard (IS).

| **Analyte** | **MRM Transitions (m/z)** | **DP (V)** | **EP (V)** | **CE (V)** | **CXP (V)** |
| --- | --- | --- | --- | --- | --- |
| GABA | 104.3 → 87.2 | 28 | 10 | 14 | 15 |
|  | 104.3 → 69.1 | 32 | 8 | 21 | 14 |
| Caffeine (IS) | 195.1 → 138.1 | 75 | 7 | 28 | 10 |
|  | 195.1 → 110.1 | 75 | 7 | 28 | 10 |

### Analysis

All electrophysiological data were analyzed using pClamp 7 and GraphPad Prism 7.0. Current density was calculated by dividing the measured peak current by the membrane capacitance (pA/pF), allowing for normalization across cells of varying size. For voltage-ramp recordings, current densities across control, glutamate, and wash conditions were compared using the Friedman test, whereas difference currents were compared using the Mann–Whitney U test. For gap-free recordings, differences in peak current density between paired conditions were assessed using the Wilcoxon signed-rank test. Normality was tested using the D’Agostino & Pearson test, and most datasets were not normally distributed. As a result, data are presented as mean values ± SEM. The normally distributed dataset was analysed using a one-way repeated measures ANOVA with Bonferroni multiple comparisons and the Geisser Greenhouse correction. All columns presented in histograms were compared with each other with a significance level of 0.05. To test whether hypoxia affected GABA release from HCs, we measured extracellular GABA concentrations ([GABA]ₑ) across several experimental conditions. [GABA]_e_ was recorded using LC-MS after cells had been incubated under hypoxic or normoxic conditions. GABA data are presented as mean ± SEM and were analyzed using a Mann-Whitney U test for pairwise comparisons.

## RESULTS

### Prolonged application of hypoxia decreased iGluR activity

Recordings were obtained from 68 HCs collected from 47 goldfish. To assess the effects of hypoxia on iGluR activity, inward Na^+^ currents evoked by iGluR activation with glutamate were recorded to measure changes in current responses to glutamate in the presence or absence of hypoxia. Current-voltage (*I-V*) relationships were produced from voltage-ramp recordings taken before and immediately after a 2 min exposure to either hypoxic or normoxic solution in the presence of 100 µM glutamate. A washout trace was included in both conditions to confirm the return to baseline following glutamate removal. In both normoxia and hypoxia, glutamate-activated currents demonstrated similar *I–V* profiles (Fig. 1A, B; *N* = 7). Glutamate application produced a significant increase in current density under both conditions, with peak current densities of −3.78 ± 1.00 pA/pF (P = 0.0010) in normoxia and −3.66 ± 0.96 pA/pF (P = 0.0323) in hypoxia at −60 mV (Fig. 1C). Difference currents for glutamate activation between normoxia and hypoxia showed a marginal reduction with mean values of −3.48 ± 0.80 pA/pF and −2.90 ± 0.74 pA/pF, respectively, though this effect was not statistically significant (Fig. 1D; P = 0.3829).

**Figure 1.**
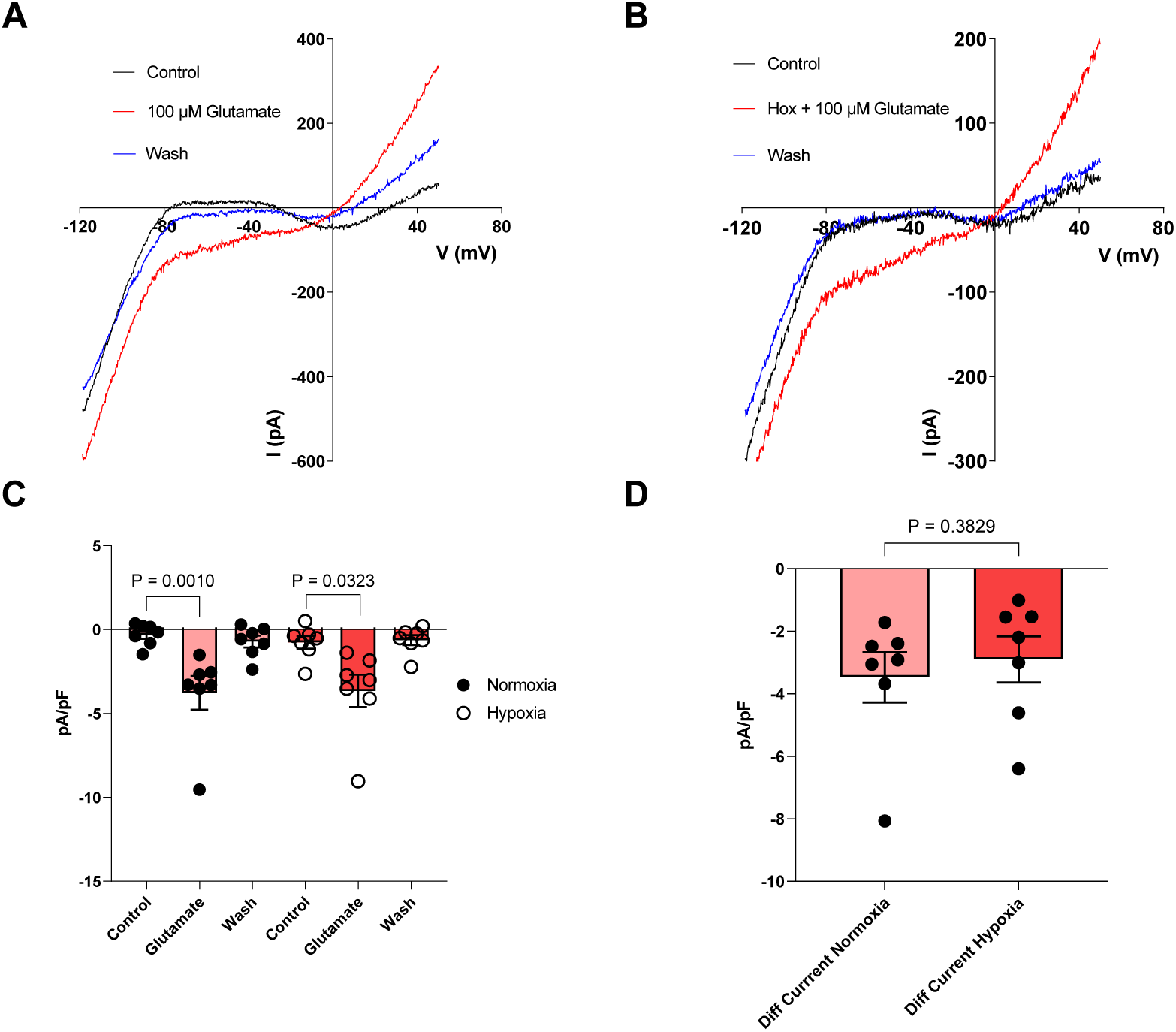
Acute hypoxia (2 min) did not affect glutamate receptor activity in goldfish horizontal cells. (**A**) Representative current-voltage (*I-V*) relationships from patch-clamp recording under normoxic conditions. Cells were exposed to extracellular solution (ECS) for 2 min, followed by application of 100 μM glutamate for 2 min, and then a 2 min washout. (**B**) Representative *I-V* relationships from a different cell in which 100 μM glutamate was applied simultaneously with hypoxia for 2 min following a 2 min washout period. Glutamate increased inward currents at negative membrane potentials under both conditions. (**C**) Summary data of current density (pA/pF) at −60 mV before, during, and after glutamate applications under normoxia (*N* = 7 from 6 fish; P = 0.0010) and hypoxia (*N* = 7 from 5 fish; P = 0.0323), showing a significant increase in normoxia and hypoxia (Friedman test). (**D**) Glutamate-activated difference currents under normoxia and hypoxia, showing no significant difference in amplitude between conditions (Mann-Whitney U test; P = 0.3829). Data are presented as mean ± SEM.

We next examined whether a prolonged application period for hypoxia would reveal a greater change in iGluR activity. Cells were clamped at −60 mV and membrane current was monitored during continuous hypoxic or normoxic exposure. Glutamate was applied twice in 2-min bouts with a washout period between successive applications. The initial and final responses to glutamate were measured from baseline to peak and compared statistically. Mean peak glutamate-activated current density decreased significantly by 24.6% after 20 min hypoxic exposure, from −4.42 ± 0.59 pA/pF during the initial bout to −3.33 ± 0.44 pA/pF during the final bout (Fig. 2A, D; *N* = 9; *P* = 0.0059). By contrast, recordings in normxia showed stable responses between glutamate applications for all cells, with no significant difference between the initial and final bouts (Fig. 2B, E; *N* = 6; P = 0.1563). Mean peak current density in normoxia was –3.61 ± 0.50 pA/pF for the first bout and –3.82 ± 0.60 pA/pF for the second.

**Figure 2.**
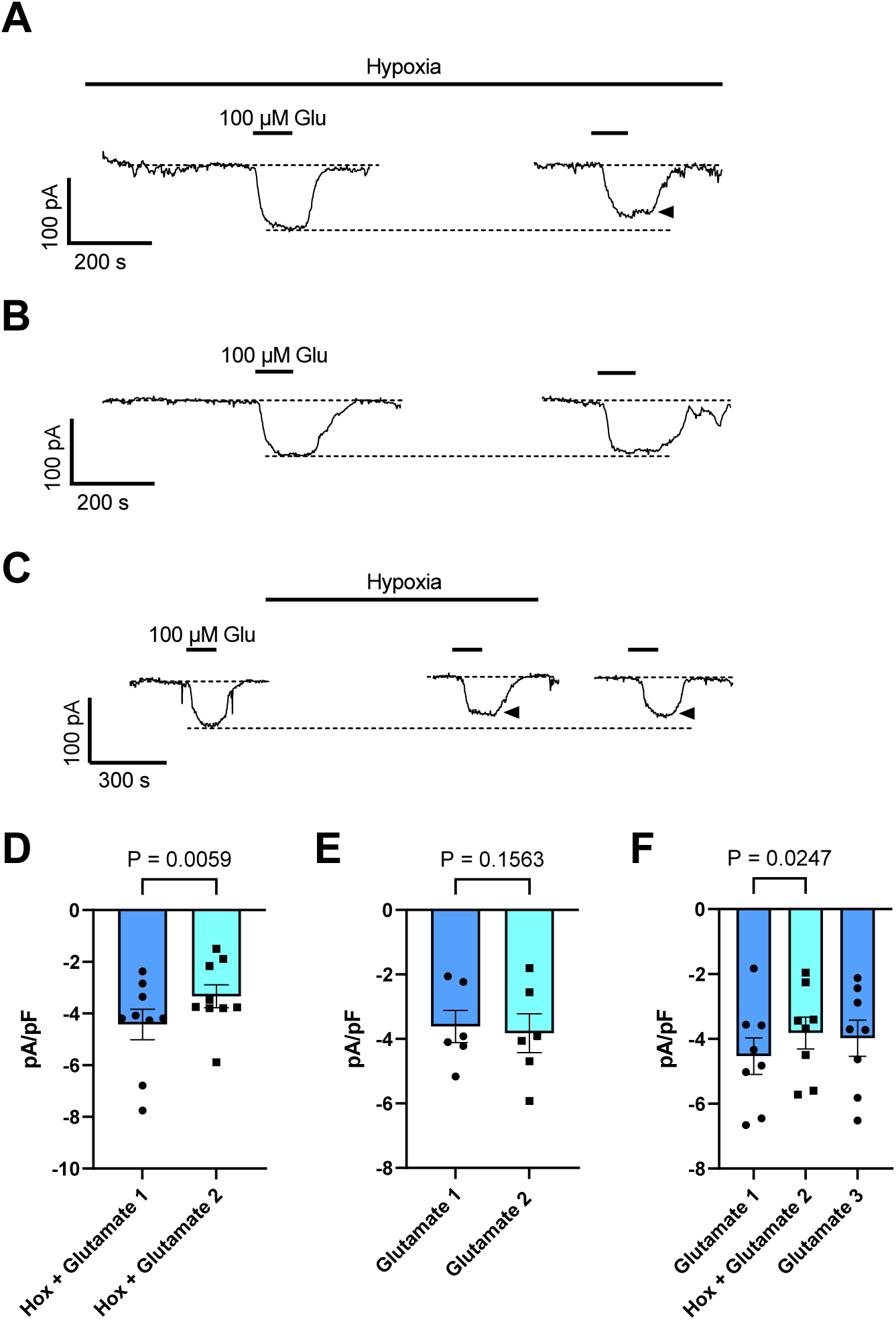
Prolonged hypoxia suppressed iGluR activity in horizontal cells. (**A**) Representative patch-clamp recording of membrane current evoked by glutamate in a horizontal cell (HC) held at −60 mV and continuously exposed to hypoxia (bar). Glutamate (100 μM) was applied for 2 min per bout. Scale bars indicate current (pA) and time (s). The second response to glutamate occurred afer 20 min hypoxia. (**B**) Representative recording from an HC clamped at - 60 mV under normoxic conditions. Glutamate (100 μM) was applied for 2 min per bout. (**C**) Representative recording from an HC voltage-clamped at −60 mV. Responses to 2 min applications of 100 μM glutamate were recorded initially (Glutamate 1), after hypoxia (Hox + Glutamate 2) and after 10 min of reoxygenation (Glutamate 3). (**D**) Summary data showing peak current density (pA/pF) for both glutamate applications under hypoxia (*N* = 9 from 8 fish), with statistical comparison between the first and second bouts showing a significant reduction in current density over time (Wilcoxon test; P = 0.0059). (**E**) Summary data under normoxia (*N* = 6 from 4 fish), showing no significant difference between the first and second bouts (Wilcoxon test; P = 0.1563). Each point represents an individual cell. (**F**) Summary data showing mean peak current density for all three bouts (*N* = 8 from 5 fish). Statistical comparisons between the first and second bout showed that hypoxia significantly lowered current density (repeated measures one-way ANOVA test; P = 0.0312) but no significant difference was measured between the second and third bouts. Each data point represents an individual cell. Each point represents an individual cell, with paired measurements obtained from the same cell across conditions. All summary data are presented as mean ± SEM.

With the hypoxia-induced suppression of glutamate-activated inward current established, we wanted to understand whether these effects were reversible. Cells were exposed to three glutamate applications, each at a specific timepoint: the first under normoxia as a control, the second after hypoxia, and the third following 10 min of reoxygenation (Fig. 2C). Under normoxia, the first glutamate application produced a strong current response with a peak current density of –4.53 ± 0.56 pA/pF on average. After hypoxic exposure, mean peak current density was reduced to –3.81 ± 0.49 pA/pF, representing a 15.9% decrease relative to the first bout (Fig. 2F; *N* = 8; P = 0.0247). Following 10 min of reoxygenation, the third glutamate response was recorded at –3.98 ± 0.56 pA/pF on average, which was not significantly different from the second bout, indicating only minor recovery. In all cells recorded, hypoxia significantly lowered iGluR responses after 15 min, but reoxygenation for 10 min did not recover current activity to control levels.

### Blocking mitochondrial KATP channels abolished the hypoxia-induced reduction in iGluR activity

To determine whether the suppression of iGluR activity during prolonged hypoxia depended on mK_ATP_ channel activation, HCs were continuously exposed to hypoxia in the presence of 100 µM glibenclamide. In the presence of glibenclamide, hypoxia did not reduce glutamate current amplitude (Fig. 3A). Summary data showed that average peak current density during the first glutamate application was −3.63 ± 0.43 pA/pF and remained stable during the second (−3.55 ± 0.39 pA/pF) application (Fig. 3B; *N* = 6). Statistical analysis revealed no significant difference in current density between bouts (P = 0.1563), indicating that blocking mK_ATP_ channels abolished the hypoxia-induced suppression of iGluR.

**Figure 3.**
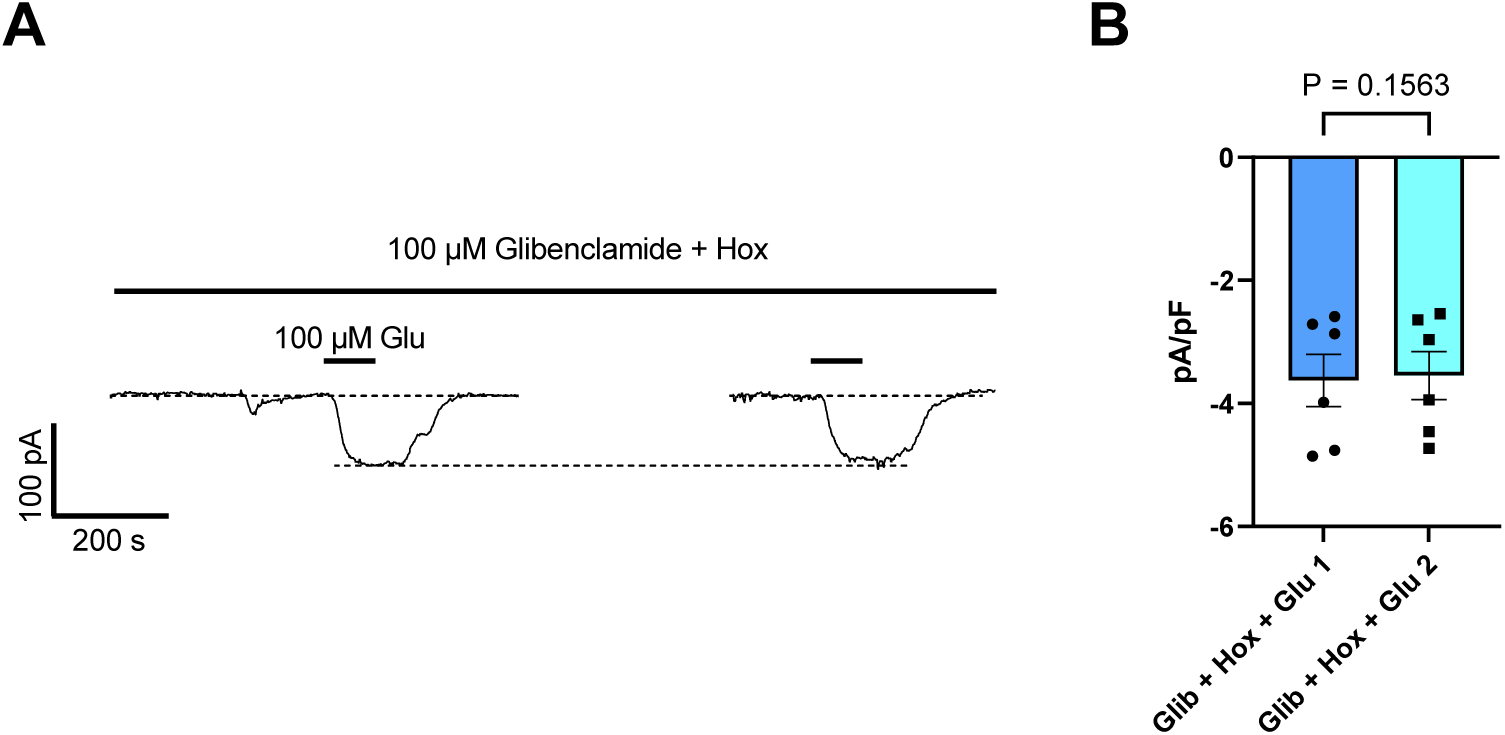
Inhibition of mK_ATP_ channels prevented hypoxia-induced suppression of iGluR. **(A)** Representative recording of membrane current in a horizontal cell clamped at −60 mV showing continuous co-application of hypoxia with 100 μM glibenclamide (bar). Two 2-min applications of 100 μM glutamate were delivered. Scale bar indicates current (pA) and time (s). **(B)** Summary data of mean peak current density (pA/pF; *N* = 6 from 4 fish). No significance was detected between the first and second application (Wilcoxon test; P = 0.1563). Inhibition of mK_ATP_ channels abolished the reduction in iGluR activity under hypoxia. Each point represents an individual cell. Data are presented as mean ± SEM.

### Hypoxia-dependent reduction in iGluR activity was inhibited by disrupting intracellular Ca^2+^ regulation

To test whether the hypoxic suppression of iGluR activity depended on intracellular Ca^2+^ signalling, we blocked Ca^2+^ uptake into the mitochondria, which would lead to an increase in [Ca^2+^]_i._ We targeted the mitochondrial Ca^2+^ uniporter (MCU) using 400 nM ruthenium red. Under hypoxia, glutamate-evoked currents remained stable across applications (Fig. 4A). Mean peak current density was −4.63 ± 1.16 pA/pF during the initial glutamate application and −4.22 ± 1.05 pA/pF for the final bout, with no significant difference between the two (Fig. 4C; P = 0.0781; *N* = 6). Blocking the MCU with ruthenium red would have elevated [Ca^2+^]_i_, thereby preventing subsequent modulation of this pathway that would otherwise occur under hypoxia.

**Figure 4.**
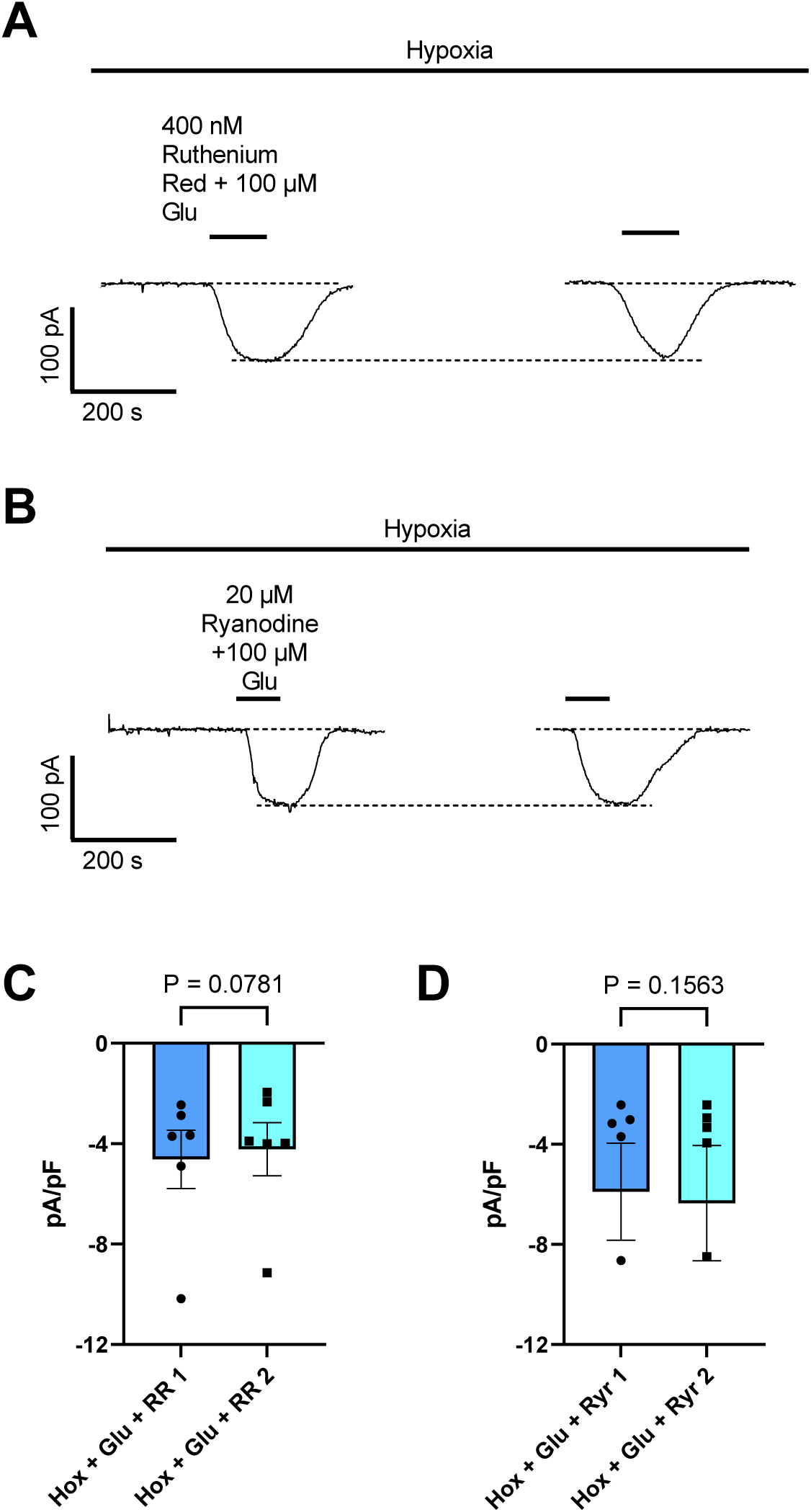
Inhibition of intracellular Ca^2+^ regulation abolished suppression of iGluR activity during hypoxia. (**A**) Representative patch-clamp recording of a horizontal cell (HC) held at −60 mV under prolonged hypoxia (bar). Ruthenium red (400 nM), used to inhibit mitochondrial Ca^2+^ uptake through the mitochondrial Ca^2+^ uniporter (MCU), was co-applied with 100 μM glutamate for 2 min per bout. Scale bars indicate current (pA) and time (s). (**B**) Representative recording of an HC voltage clamped at −60 mV under prolonged hypoxia. Ryanodine at 20 μM, used to inhibit Ca^2+^ release through ryanodine receptors in the endoplasmic reticulum, was co-applied with 100 μM glutamate for 2 min per bout. (**C**) Summary data for ruthenium red treatment, showing no significant difference in peak current density (pA/pF) between the glutamate applications (*N* = 6 from 5 fish; Wilcoxon test; P = 0.0781). (**D**) Summary data for ryanodine treatment, showing no significant difference in peak current density between the glutamate applications (*N* = 6 from 3 fish; Wilcoxon test; P = 0.1563). Each point represents a measurement from an individual cell, with paired measurements obtained from the same cells during applications. Data are presented as mean ± SEM.

Next, we blocked Ca^2+^ release from the ER by applying 20 µM ryanodine. Current amplitudes remained stable over time (Fig. 4B). Mean peak current density was −5.89 ± 1.94 pA/pF in the first bout and −6.35 ± 2.31 pA/pF in the second, indicating that blocking Ca^2+^ release through ryanodine receptors abolishes the hypoxia-induced suppression in the presence of glutamate (Fig. 4D; P = 0.1563; *N* = 6).

### Inhibition of calmodulin and phosphatases prevented the hypoxia-induced reduction in iGluR activity

To determine whether the Ca^2+^-dependent suppression of iGluR activity involved downstream calmodulin and phosphatase signalling, both pathways were examined pharmacologically. Calmodulin is an intracellular Ca^2+^ binding protein that translates changes in [Ca^2+^]_i_ into downstream signalling, whereas protein phosphatase 1 and 2A (PP1/PP2A) regulate iGluR activity through receptor dephosphorylation. Both pathways have been demonstrated in the reversible suppression of glutamate receptor current during anoxia in turtle neurons (Shin et al., 2005; Pamenter et al., 2008; reviewed in Buck & Pamenter, 2018; Bickler et al., 2000). HCs were treated with 15 µM calmidazolium, a calmodulin inhibitor. Under hypoxia, glutamate-evoked currents remained stable across application (Fig. 5A). Mean peak current density was - 5.51 ± 1.48 pA/pF during the first glutamate application and −5.10 ± 1.16 pA/pF during the second application, with no significant difference between the two responses (Fig. 5C; P = 0.4063; *N* = 7).

**Figure 5.**
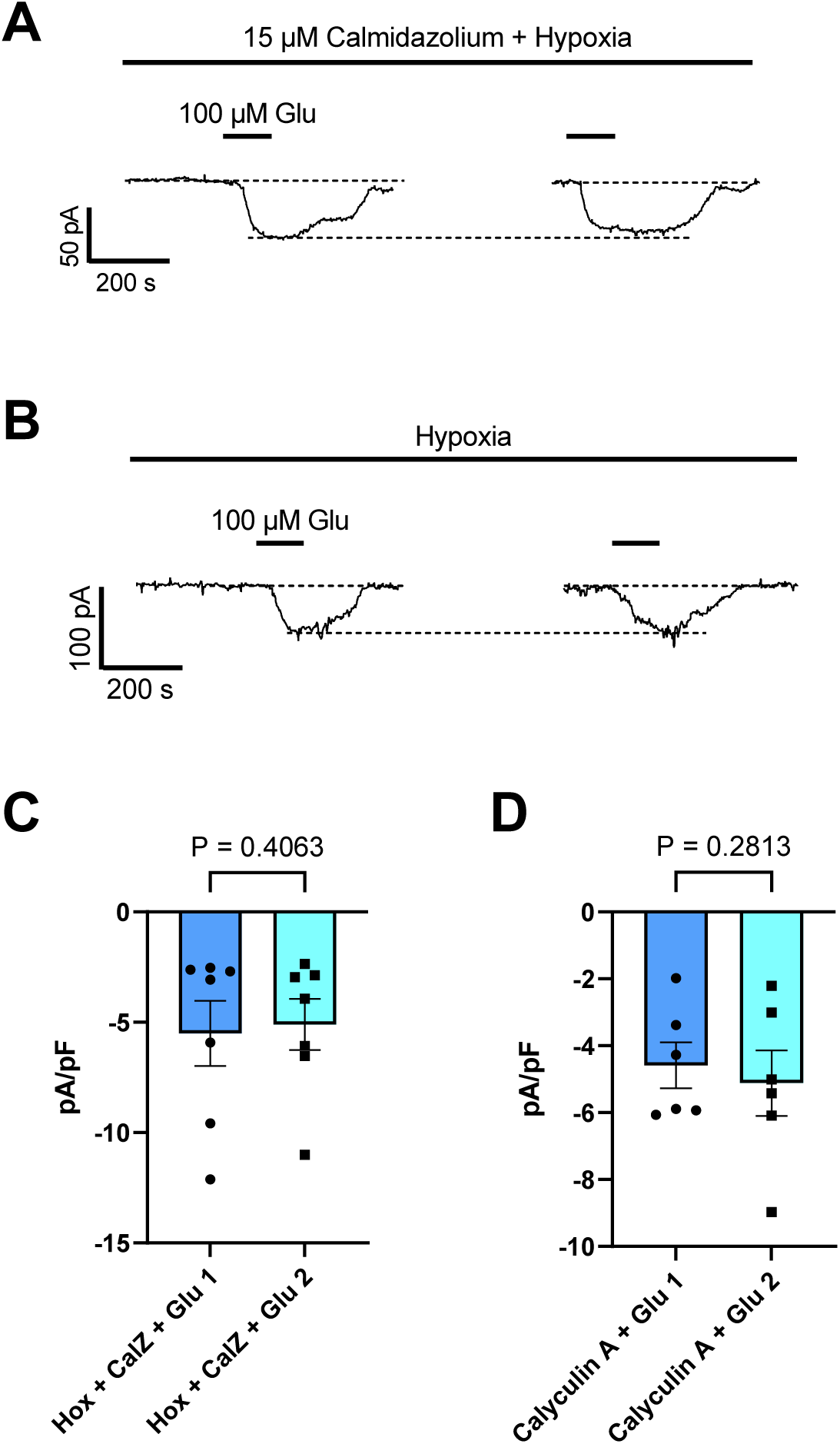
Effects of calmodulin and protein phosphatase inhibition on iGluR during prolonged hypoxia. (**A**) Representative recording of membrane current in a horizontal cell (HC) held at −60 mV under continuous hypoxia. The calmodulin inhibitor, calmidazolium, was continuously applied at 15 μM (bar). Glutamate at 100 μM was applied for 2 min per bout and responses to glutamate are shown. Scale bars indicate current (pA) and time (s). (**B**) Representative recording from an HC held at −60 mV under continuous hypoxia. The cell was pre-incubated for 30 min with 10 nM calyculin A, an inhibitor of protein phosphatases 1 and 2A (PP1/PP2A). Glutamate at 100 μM was applied for 2 min per bout. (**C**) Summary data for calmidazolium treatment, showing no significant difference in peak current density (pA/F) between glutamate applications (*N* = 7 from 4 fish; Wilcoxon test; P = 0.4063). (**D**) Summary data for calyculin A treatment showing no significant difference in peak current density between glutamate applications (*N* = 6 from 3 fish; Wilcoxon test; P = 0.2813). Each point represents a measurement from an individual cell, with paired measurements obtained from the same cells across the displayed applications. Data are presented as mean ± SEM.

To determine whether protein phosphatase (PP1/PP2A) activity contributed to this inhibitory pathway, cells were preincubated with 10 nM calyculin A for 30 min before recordings were performed. Glutamate-evoked currents again remained stable during hypoxic exposure (Fig. 5B). Mean peak current density was −4.59 ± 0.69 pA/pF during the initial bout of glutamate and −5.12 ± 0.98 pA/pF during the final bout, with no significant difference observed (Fig. 5D; P = 0.2813; *N* = 6). Thus, inhibition of PP1/PP2A pathways abolished the hypoxia-induced suppression of iGluR activity.

### Increased [Ca^2+^]i did not contribute to GABA release during hypoxia

An alternative potential outcome of elevated [Ca^2+^]_i_ during hypoxia is that Ca^2+^-dependent secretion of GABA will be increased. Extracellular GABA concentration ([GABA]ₑ) was measured across several experimental conditions using LC-MS after cells had been incubated under hypoxic or normoxic conditions. An overview of this experimental procedure is shown in Figure 6. In control dishes [GABA]ₑ was 78.66 ± 7.49 nM, whereas in dishes treated with glutamate [GABA]ₑ was significantly greater at 552.90 ± 241.7 nM (Fig. 7A; P = 0.0500; N = 6). These findings confirm sensitive detection of GABA release from depolarized HCs using LC-MS. A chromatogram of glutamate-induced GABA release is provided in Figure 7B. Both traces correspond to the same sample, with one MRM signal used for GABA semi-quantification (black trace) and the other used to confirm GABA peak identity (grey trace).

**Figure 6.**
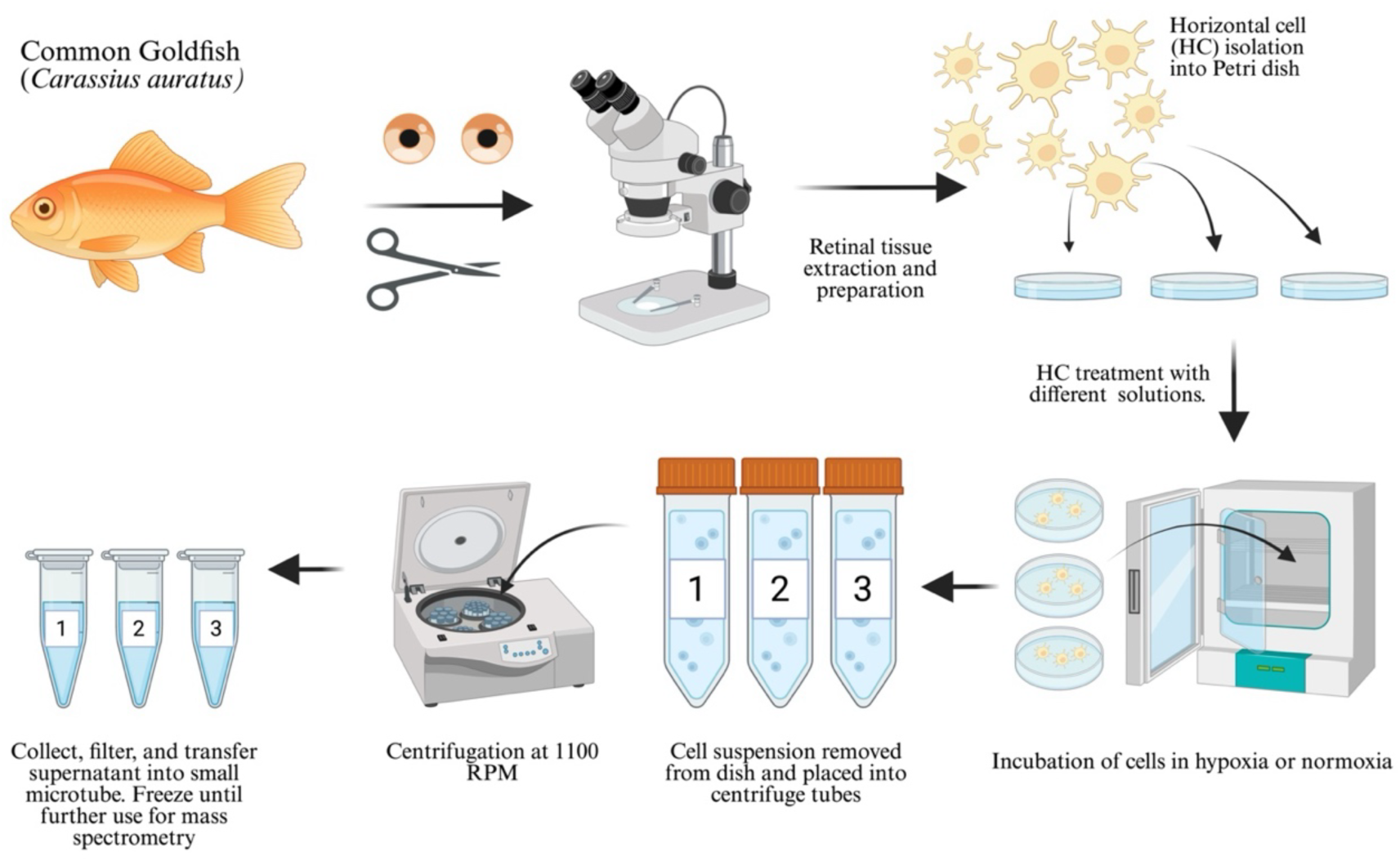
Overview of protocol to detect GABA release from goldfish horizontal cells by LC-MS. From the upper-left moving clockwise, retinas were removed from goldfish and horizontal cells were dissociated *in vitro*. Tissue from both retinas of each fish were divided into three equal suspensions and placed in Petri dishes. Dishes were placed in an incubator and exposed to glutamate, hypoxia (PO_2_ = 15 mmHg), hypoxia + glutamate, or to extracellular solution as control. After 1-h exposures, cells were transferred to large centrifuge tubes (1-3) and centrifuged. The cell-free supernatant was collected, filtered, and stored at –20 °C in microtubes for subsequent quantification of γ-aminobutyric acid (GABA) concentration by LC-MS (see Methods for details).

**Figure 7.**
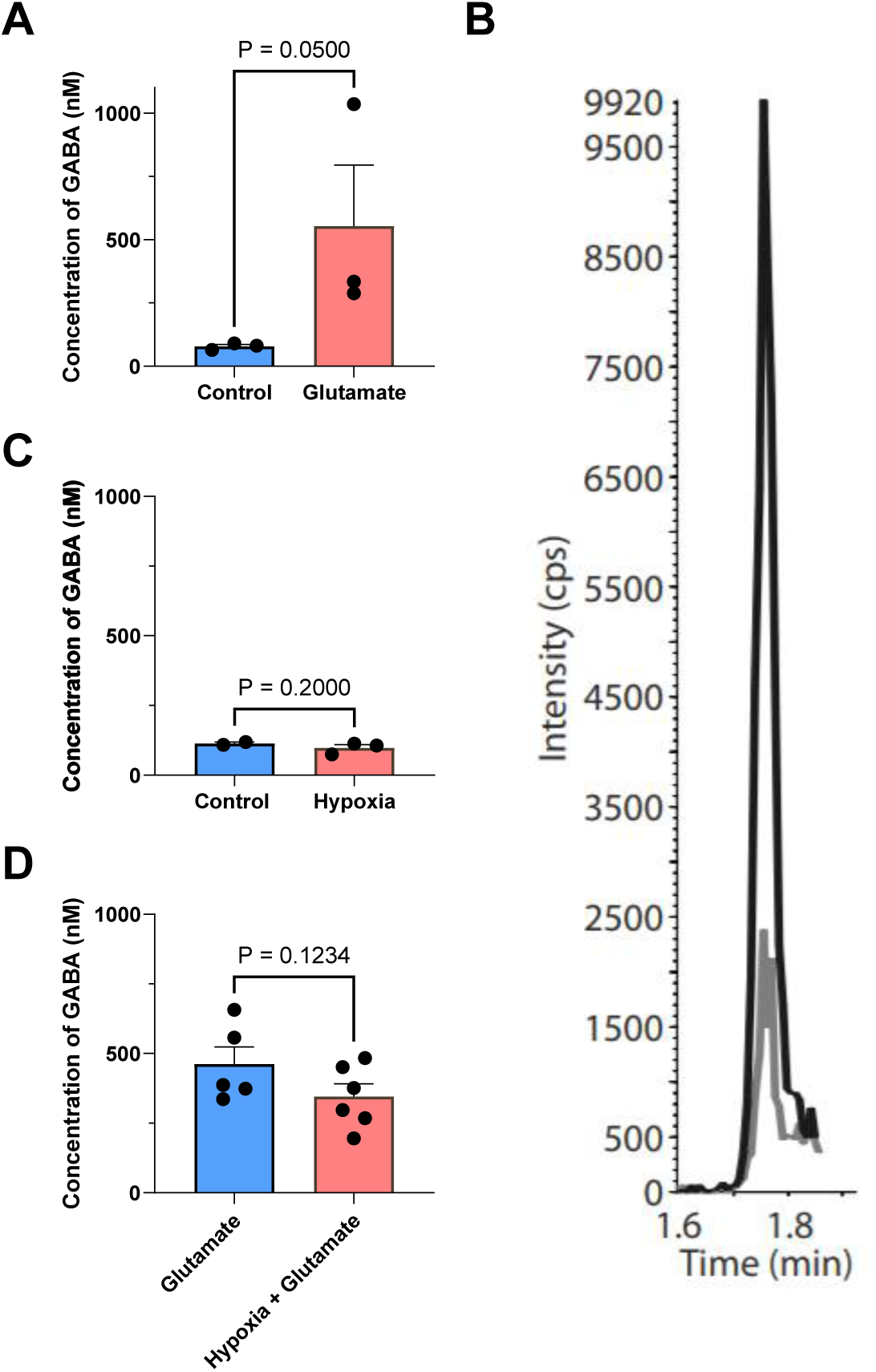
Hypoxia did not increase GABA release from goldfish horizontal cells. (**A**) Concentration of extracellular γ-aminobutyric acid ([GABA]_e_) released from isolated horizontal cells (HCs) following a 1-h incubation period in 100 μM glutamate, measured by LC-MS. A significantly higher [GABA]_e_ was induced by glutamate, compared to untreated controls (*N* = 6 from 2 fish; Mann Whitney U; P =0.05). (**B**) Representative LC-MS chromatogram of a single sample from the glutamate-treated group in (A). Multiple reaction monitoring (MRM) signals from the same sample were used for GABA semi-quantitation (black trace) and confirmation of GABA peak identity (grey trace). The y-axis represents intensity (counts per second, cps) and the x-axis shows the retention time (min). (**C**) [GABA]_e_ in cells incubated for 1 h under control (normoxia) or hypoxia did not show any significant difference (*N* = 5 from 2 fish; Mann-Whitney U test; P = 0.2000). (**D**) [GABA]_e_ after 1 h incubation in glutamate or hypoxia co-applied with glutamate. GABA release did not differ between hypoxic and normoxic treatment conditions (*N* = 11 from 4 fish; Mann-Whitney U test; P = 0.1234).

To assess the effect of hypoxia alone on GABA release, cells were incubated under normoxic or hypoxic conditions for 1 h. Mean [GABA]_e_ was 113.80 ± 4.85 nM in normoxia and 97.85 ± 11.90 nM under hypoxia, with no significant difference between treatments (Fig. 7C; *N* = 5; *P* = 0.2000). Co-application of glutamate with hypoxia yielded a mean [GABA]_e_ of 345.40 ± 45.56 nM, which was not significantly different compared to [GABA]_e_ from cells treated with glutamate under normoxic conditions (462.10 ± 61.77 nM; Fig. 7D; P = 0.1234; *N* = 11).

## DISCUSSION

### iGluR suppression during hypoxia

The present study demonstrates that HCs isolated from goldfish retina display a decrease in glutamate-activated currents following exposure to prolonged (20 min) hypoxia. This response involves the activation mK_ATP_ channels and the subsequent release of intracellular Ca^2+^ from stores in the ER, linking mitochondrial signalling to suppression of membrane iGluR activity.

Our findings further show that this response is dependent on calmodulin and PP1/PP2A activity, supporting the involvement of Ca^2+^-sensitive phosphatase signalling in the hypoxia-induced reduction of iGluR. Building on previous studies from our lab, these results support a model, where hypoxia activates mK_ATP_ channels in goldfish HCs. In the absence of glutamate, mK_ATP_ activation maintains [Ca^2+^]_i_ (Country & Jonz, 2021); whereas in the presence of extracellular glutamate, mK_ATP_ activation during hypoxia paradoxically triggers a modest release of Ca^2+^ from internal stores (Nagy-Watson & Jonz, 2025; reviewed in Jonz, 2025).

Much of the ATP consumption in the retina is used to maintain ionic gradients across the plasma membrane, which is essential for neural signalling (Wong-Riley 2010). When activated, iGluRs are permeable to Na^+^ and Ca^2+^ (Dingledine et al., 1999; Diamond, 2011), which causes depolarization and opens voltage-gated Ca^2+^ channels. The influx of ions ultimately requires the work of the Na^+^/K^+^-ATPase, and other ion transport mechanisms, to restore the cell to resting membrane potential. Thus, each cycle of depolarization and ionic gradient restoration requires a significant amount of energy to maintain, making the retina particularly sensitive to changes in oxygen availability. In hypoxia-tolerant vertebrates, one strategy to reduce this burden is by actively downregulating receptor activity, thereby decreasing ion flux and lowering ATP demand (Pamenter et al., 2008; Buck & Bickler, 1998; Wilkie et al., 2008; Bickler et al., 2000; reviewed in Buck & Pamenter, 2018). In crucian carp (*Carassius carassius*), a close relative to the goldfish, responses to light in both the retina and optic tectum are reversibly suppressed during prolonged anoxia. Specifically, response amplitudes decreased by 69% and 75% respectively after 38 min of anoxia and by 90% after 1 h (Johansson et al., 1997). Similarly, Wei & Yang (1997) reported that cone- and rod-driven HCs in crucian carp show a reduction in response amplitude to 76% and 55%, respectively, from the controls after 10 min of hypoxia. Together, these findings support the idea that iGluR activity in HCs is downregulated during hypoxia and that this suppression develops gradually, rather than occurring immediately when oxygen levels drop.

The present study reports no significant difference in glutamate-evoked current amplitude during acute hypoxia (2 min). By contrast, prolonged hypoxia resulted in a clear reduction in glutamate responses. Mean peak current density declined by ∼25% after 20 min of hypoxic exposure. This effect was absent under normoxia, where responses remained stable across both glutamate applications. The stability of recordings during normoxia indicate that receptor desensitization or cell rundown did not contribute to the observed iGluR suppression, suggesting that hypoxia alone induced the reduction in receptor activity at the plasma membrane.

We found that hypoxic suppression of iGluR activity did not recover following up to 10 min of recovery in normoxic recording solution. Due to technical limitations in maintaining stable patch-clamp recordings for extended durations, we were unable to assess recovery beyond that time. However, this lack of recovery should not be interpreted as evidence of irreversibility. Rather, the recovery process may require a longer reoxygenation period. Indeed, in other hypoxia-tolerant vertebrate systems, recovery from hypoxic suppression often requires a reoxygenation period longer than 10 min. In crucian carp, supressed responses in the retina required approximately 45 min of normoxia to recover from a 90% suppression (Johansson et al., 1997). Similarly, in hippocampal slices of the naked mole rat, synaptic activity remained supressed after 30 min of anoxia and recovered after 30-60 min reoxygenation periods (Larson & Park, 2009). In the anoxia-tolerant turtle, AMPA receptor currents returned to control levels after approximately 40 min of reoxygenation, while recovery of cortical EEG activity following prolonged anoxia required nearly 100 min of reoxygenation (Pamenter et al., 2008; Fernandes et al., 1997).

Additional studies have shown that recovery of neural responses to baseline excitability following anoxia can require prolonged periods of restored oxygen availability, likely due to the time needed to reestablish ionic gradients and recover from metabolic stress (Money et al., 2014; Edelman et al., 1991). It is therefore possible that goldfish HCs require lengthy reoxygenation periods as well to demonstrate significant recovery from the hypoxia-induced suppression of iGluRs. Future work extending reoxygenation periods could provide further insight into the potential reversibility of this response.

### mKATP channel activation as a sensor and mediator of receptor downregulation

In the current study, the decrease in iGluR activity during hypoxia depended on the activation of mK_ATP_ channels, as blocking these channels with glibenclamide eliminated the suppression response. In goldfish HCs, fluorescence of tetramethylrhodamine ethyl ester (TMRE) dye was used to monitor changes in mitochondrial membrane potential (Δψ_m_). TMRE fluorescence increased significantly within the first 5 min of hypoxia, but this response was reduced in the presence of mK_ATP_ antagonists, like 5-hydroxydecanoate (5-HD) and glibenclamide. These findings indicated that mK_ATP_ channels are activated during the first few minutes of hypoxia and could be functioning as metabolic sensors (Country & Jonz, 2021) during low oxygen periods to trigger a neuroprotective response.

In several animal models, activation of mK_ATP_ channels during hypoxia serves as the initiating signal for neuroprotection. In the mammalian retina, brief ischemic or hypoxic preconditioning has been shown to protect neurons against subsequent injury, with the opening of mK_ATP_ channels playing a central role in this response (Yamauchi et al., 2003; Roth, 2004; Roth et al., 2006). Similarly, in primary rat cortical neurons, opening of mK_ATP_ channels preconditioned cells against glutamate excitotoxicity and oxygen deprivation (Kis et al., 2004). In the anoxia-tolerant turtle, mK_ATP_ activation suppressed AMPA receptor currents by 52% during anoxia; and activation of mK_ATP_ channels during normoxia produced a similar decrease in AMPA receptor activity (Zivkovic & Buck, 2010). In the hypoxia-intolerant rainbow trout, activation of mK_ATP_ with diazoxide maintained [Ca^2+^]_i_ during hypoxia, whereas without activation, [Ca^2+^]_i_ rose to excitotoxic levels within 20 min (Country & Jonz, 2021).

When mK_ATP_ channels open during anoxia, Δψ_m_ slightly depolarizes due to the partial dissipation of the mitochondrial H^+^ gradient. This results in a modest increase in cytosolic Ca^2+^ due to a reduction in the driving force of the MCU, which limits Ca^2+^ uptake into the mitochondria, as well as increases Ca^2+^ release from the mitochondrial membrane transition pore (MPTP) (Pamenter, 2014; Buck & Pamenter, 2018; Hawrysh & Buck, 2013). In HCs, Ca^2+^-induced Ca^2+^ release (CICR) also plays an important role in regulating Ca^2+^ dynamics (reviewed in Country, 2017). Calcium imaging experiments in the carp retina have shown that, in the presence of glutamate, there is a large transient peak in [Ca^2+^]_i_ that is blocked by ryanodine, suggesting it is released from CICR through ER stores (reviewed in Country, 2017; Huang et al., 2004). Thus, it was important to determine whether cytosolic Ca^2+^ was mediating the suppressed response during hypoxia.

### A potential role for intracellular Ca^2+^ in hypoxia tolerance

As predicted, blocking Ca^2+^ uptake via the MCU prevented the suppression of iGluR current under hypoxia, suggesting that mitochondrial cytosolic Ca^2+^ buffering contributes to this response. When the MCU was inhibited with 400 nM ruthenium red, glutamate responses remained stable throughout hypoxia. Similarly, blocking Ca^2+^ release from the ER with 20 μM ryanodine produced the same outcome. Although inhibition of the MCU would reduce mitochondrial Ca^2+^ buffering, and potentially increase cytosolic Ca^2+^, uptake through the MCU under hypoxia may potentially serve to maintain [Ca^2+^]_i_ at levels that evade excitotoxicity (Nicholls & Budd, 2000; Rizzuto et al., 2012) while also contributing to the regulated Ca^2+^ signal. These findings support those of Nagy-Watson & Jonz (2025) who found that blocking the MCU or ryanodine receptors in the ER abolished a hypoxia-induced rise in intracellular Ca^2+^ in goldfish HCs, supporting the idea that both the mitochondria and ER contribute to initiating the neuroprotective suppression of iGluR activity. Together these results indicate that regulation of mitochondrial Ca^2+^ uptake and Ca^2+^ release from the ER may be required to initiate the neuroprotective suppression of iGluR activity.

The downstream components of the Ca^2+^-dependent pathway were also investigated by inhibiting calmodulin and other protein phosphatases. Treatment with 15 μM calmidazolium prevented the hypoxia-induced suppression of glutamate-evoked currents, indicating that calmodulin signalling contributes to this response. Calmidazolium is a potent, membrane permeable calmodulin antagonist, and 15 μM has previously been used to inhibit calmodulin-dependent signalling in intact cells and neuronal electrophysiological preparations (Chang et al., 2000; Houchat et al., 2022). Similarly, treatment with 10 nM calyculin A for 30 min prevented the reduction in iGluR activity during hypoxia, indicating that activity of a calyculin A-sensitive phosphatase is required for this response. Calyculin A is a potent inhibitor of both PP1 and PP2A. In intact MCF-7 cells, exposure to 10 nM calyculin A inhibited the activities of both phosphatases by more than 50% (Favre et al., 1997). Comparable treatment conditions have been used to investigate phosphatase-dependent channel regulation, with cells exposed to 10 nM calyculin A for 10-30 min before patch clamp recordings (Sansom et al., 1997) and for 30 min to alter intracellular protein phosphorylation in HEK292T cells (Grabinski & Kanaan, 2016). PP1-dependent regulation of iGluR has been previously demonstrated (Westphal et al., 1999; Yan et al., 1999), while Ca^2+^/calmodulin can directly promote NMDA receptor inactivation (Ehlers et al., 1996). These findings are consistent with the involvement of calmodulin signalling and PP1/PP2A activity in the pathway linking intracellular Ca^2+^ signalling to the suppression of iGluR activity during hypoxia.

This rise in [Ca^2+^]_i_ may engage calmodulin and phosphatase-dependent signaling pathways that ultimately reduce iGluR activity at the plasma membrane (Bickler et al., 2000; Shin et al., 2005). The precise relationship between calmodulin and PP1/PP2A, including whether they act sequentially or through parallel pathways, remains to be established. In the present model, in which glutamate is present and the membrane depolarized, we propose that hypoxia leads to activation of mK_ATP_ (Country & Jonz, 2021). Paradoxically, Ca^2+^ is released from internal stores, resulting in a modest yet significant rise in [Ca^2+^]_i_ (Nagy-Watson & Jonz, 2025). The effects of calmodulin are best characterized for NMDA receptors. Ca^2+^-bound calmodulin binds directly to the intracellular C-terminal region of the GluN1 subunit, reducing channel open probability and producing Ca^2+^-dependent inactivation. Calmodulin also competes with α-actinin for binding to GluN1, weakening the interaction between the receptor and the actin cytoskeleton, and potentially facilitating its removal from the membrane (Ehlers et al., 1996; Zhang et al., 1998). In parallel, PP1 can be anchored to NMDA receptors through scaffold proteins, where its phosphatase activity limits receptor currents (Westphal et al., 1997). PP2A can also associate with NMDA receptor subunits and regulate receptor phosphorylation, providing an additional mechanism through which phosphatase activity may reduce receptor function (Chan & Sucher, 2001). In turtle cortical neurons, inhibition of both calmodulin or PP1/PP2A prevents the anoxia-induced reduction in NMDA receptor currents, indicating that both components contribute to channel arrest (Shin et al., 2005). Although this mechanism was initially described for NMDA receptors, Ca^2+^-permeable AMPA receptors are the predominant iGluR subtype in *Carassius* HCs and likely mediate the currents recorded here (Sun et al., 2010). AMPA receptor activity is also regulated through receptor phosphorylation and trafficking, and AMPA receptor currents undergo reversible, mK_ATP_-dependent suppression in anoxic turtle neurons (Banke et al., 2000; Snyder et al., 2003; Pamenter et al., 2008a; Zivkovic and Buck, 2010). Therefore, a similar pathway may supress AMPA receptor activity in goldfish HCs during hypoxia.

The proposed pathway is similar to a mechanism described in the cortex of the anoxia-tolerant turtle brain. The modest rise in cytosolic Ca^2+^ acts as a signalling cue that interacts with calmodulin and PP1/PP2A, which in turn inhibits iGluRs and promotes channel arrest (Pamenter et al., 2008; reviewed in Buck & Pamenter, 2018; Shin et al., 2005; Ehlers et al., 1996; Krupp et al., 1999; Chowdhury et al., 2017). Complementary work in the same species demonstrated that AMPA receptors are also downregulated during anoxia (Pamenter et al., 2007). Together, these findings suggest that a similar pathway may occur in goldfish HCs, where AMPA receptor suppression could be the primary mechanism for the observed decrease in glutamate-activated currents, ultimately conserving energy and reducing Ca^2+^ influx during hypoxia. This mechanism is summarized in Figure 8 and is supported by the work of Johansson et al. (1997), who demonstrated that retinal responses are strongly suppressed during anoxia in crucian carp. In this context, the suppression of iGluR activity in HCs would decrease excitability in HCs, potentially contributing to overall reduction in retinal activity. The presented mechanism may also promote glutamate release from photoreceptors. Suppression of iGluR activity would hyperpolarize HCs, which would reduce proton-mediated inhibition of presynaptic Ca^2+^ channels. The resulting increase in presynaptic Ca^2+^ influx would increase glutamate release from photoreceptors, which in turn would hyperpolarize ON bipolar cells. Whether this response depends solely on HCs as hypoxia sensors or also involves other retinal neurons remains to be determined.

**Figure 8.**
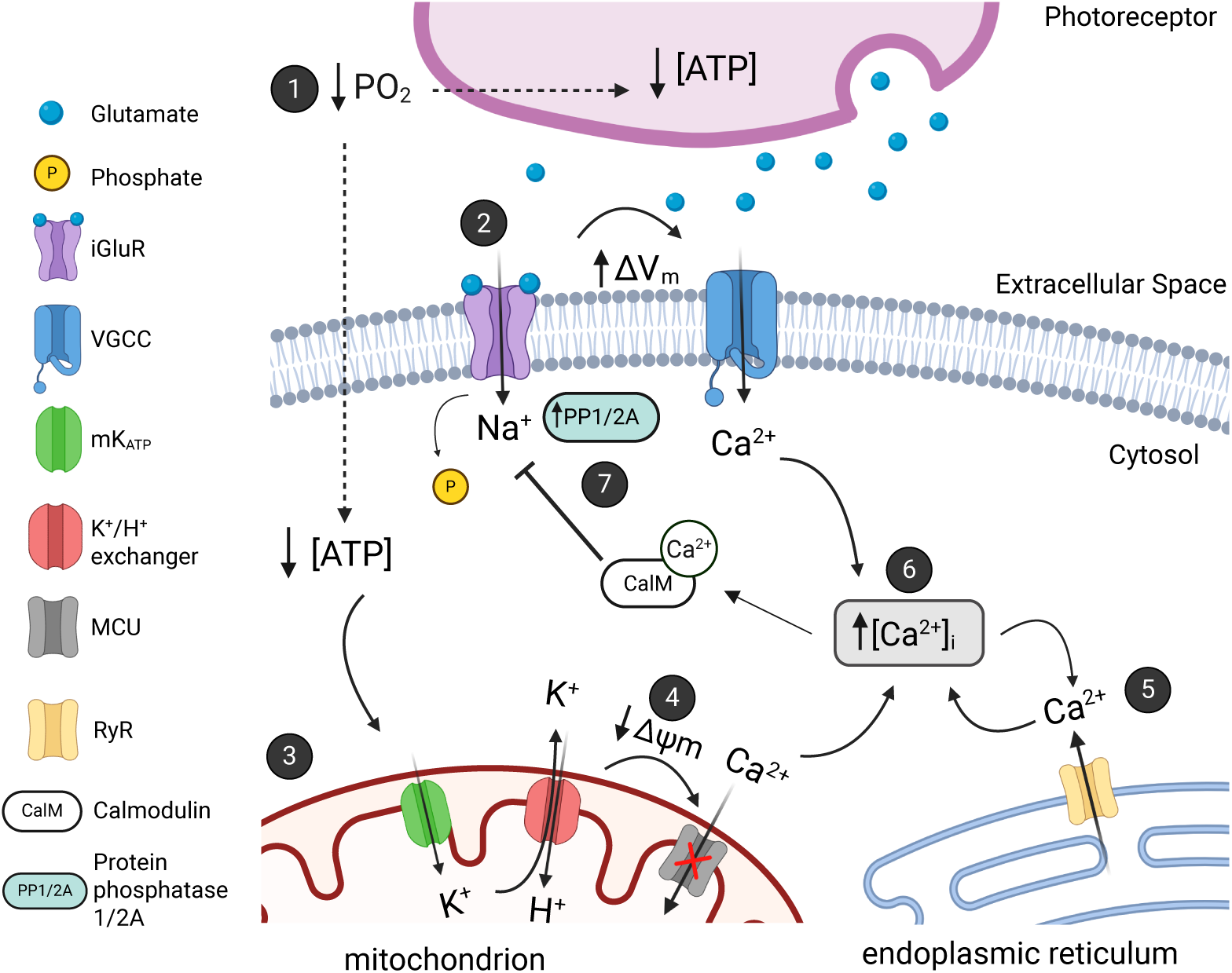
Working model of a hypoxia-tolerance mechanism through which intracellular Ca^2+^ signalling induces the downregulation of ionotropic glutamate receptors in horizontal cells of goldfish retina. Starting at the upper left, hypoxia (↓PO_2_) leads to a decline in ATP levels in both photoreceptors and HCs, as reduced oxygen availability limits ATP production through oxidative phosphorylation (1). In photoreceptors, this decrease in ATP availability leads to uncontrolled depolarization and excessive glutamate release, which activates ionotropic glutamate receptors (iGluR, purple) on the horizontal cell (HC) membrane. Binding of glutamate (2) increases inward Na^+^ current, resulting in the depolarization of the membrane (ΔV_m_) and subsequent opening of voltage gated Ca^2+^ channels (VGCC, blue). In HCs, hypoxia also reduces ATP production, which lowers the ATP:ADP ratio and thereby triggers the opening of mitochondrial ATP-dependent K^+^ channels (mK_ATP_, green) on the inner mitochondrial membrane (Tinker et al., 2014) (3). The activation of these channels increases K^+^ influx across the mitochondrial membrane, which leads to ion exchange (K^+^/H^+^ exchanger, red) and produces a modest depolarization of the mitochondrial membrane potential (Δψm) (4). This reduces the uptake of Ca^2+^ via the mitochondrial Ca²⁺ uniporter (MCU, grey), allowing cytosolic Ca^2+^ levels to rise. Together with Ca^2+^ entry through VGCCs in the membrane, this signal promotes Ca²⁺-induced Ca²⁺ release from endoplasmic reticulum (ER) via ryanodine receptors (RyR, yellow) (5), which ultimately further increases [Ca²⁺]_i_ (6). Elevated [Ca²⁺]_i_ activates calmodulin (CalM), which may bind to iGluR or associated proteins to reduce receptor activity. Protein phosphatases 1 and 2A (PP1/PP2A) may also contribute through receptor dephosphorylation (7). This model is supported by evidence from goldfish HCs (Country & Jonz, 2021; Nagy-Watson & Jonz, 2025; reviewed in Jonz, 2025) as well as a similar mechanism described in the anoxia-tolerant turtle brain (Buck & Pamenter, 2018; Pamenter et al., 2008). Through this mechanism, mK_ATP_ activation and subsequent Ca^2+^ signalling work together to reduce excitotoxic glutamatergic responses during hypoxia by conserving energy and limiting Ca^2+^ overload. For further details, see text. Created using BioRender.com

### Glutamate-induced GABA release was insensitive to hypoxia

As an important site of the storage and release of GABA in the outer retina (Ayoub and Lam, 1984,1985; Barnes et al., 2020; Marc et al., 1978), HCs were proposed to induce retinal suppression via GABA secretion (Johansson et al., 1997). To assess whether the rise in [Ca^2+^]_i_ during hypoxia might initiate vesicular release of GABA, we used LC-MS to quantify [GABA]_e_ and found that hypoxia did not change GABA secretion when HCs were depolarized by glutamate. These results argue against the idea that Ca^2+^ plays a role in increasing GABA release onto photoreceptors during hypoxia to supress retinal activity.

## CONCLUSION

The present study demonstrates that iGluR activity in goldfish HCs is significantly reduced during prolonged hypoxia, and that this suppression is dependent on the activation of mK_ATP_ channels. Activation of these channels initiates a controlled rise in intracellular Ca^2+^ from internal stores (Nagy-Watson & Jonz, 2025), which serves as a signal to downregulate iGluR activity, potentially preventing further excitotoxic Ca^2+^ accumulation (Country & Jonz, 2021). A similar Ca^2+^-dependent suppression pathway has been described in the anoxia-tolerant turtle cortex (Pamenter et al., 2008), suggesting a conserved neuroprotective strategy among vertebrates under metabolic stress. These findings provide insight into a potential mechanism in goldfish retina, where receptor activity in the membrane is modulated to reduce metabolic demand, maintain [Ca^2+^]_i_ and preserve cell viability during stress. Taken together, these results suggest that HCs act as sensors of hypoxia in goldfish, initiating a response that maintains neuronal function, and may underlie suppression of retinal activity in hypoxia-tolerant animals. Whether other retinal neurons also exhibit a similar sensitivity to oxygen availability remains an open question and will require further investigation. Understanding the mechanisms that enable retinal neurons of hypoxia-tolerant vertebrates to survive metabolic stress may reveal potential targets for promoting retinal neuroprotection in all vertebrates, include those that cannot withstand hypoxia. These mechanisms may be relevant to developing interventions against retinal diseases, many of which involve hypoxia or ischemia (Osborne et al., 2004), and to emerging cell-based therapies, such as where metabolic stress and mitochondrial dysfunction contribute to early donor cell loss following retinal transplantation (Sudharsan et al., 2025).

## Acknowledgements

The authors thank Dr. Christian A. Rosales and Dr. Jeffrey C. Smith of the Carleton Mass Spectrometry Centre at Carleton University for performing liquid chromatography-mass spectrometry analysis of GABA samples.

## Author contributions

Conceptualization: M.G.J., J.C.S.; Methodology: all authors; Validation: all authors.; Formal analysis: M.G.J., M.R.; Investigation: all authors.; Resources: M.G.J., J.C.S.; Data curation: M.R.; C.A.R.; Writing - original draft: all authors; Writing - review & editing: all authors; Supervision: M.G.J., J.C.S.; Project administration: M.G.J.; Funding acquisition: M.G.J., J.C.S.

## Data availability

All data associated with this work are presented in the manuscript.

